# Identification of novel risk loci with shared effects on alcoholism, heroin and methamphetamine dependence

**DOI:** 10.1101/505917

**Authors:** Yan Sun, Suhua Chang, Zhen Liu, Libo Zhang, Fan Wang, Weihua Yue, Hongqiang Sun, Zhaojun Ni, Xiangwen Chang, Yibin Zhang, Yang Chen, Jiqiang Liu, Lin Lu, Jie Shi

## Abstract

Different substance dependences have common effects on reward pathway and molecular adaptations, however little is known regarding their shared genetic factors. We aimed to identify the risk genetic variants that are shared for substance dependence (SD). First, promising genome-wide significant loci were identified from 3296 patients (521 alcoholic/1026 heroin/1749 methamphetamine) vs 2859 healthy controls and independently replicated using 1954 patients vs 1904 controls. Second, the functional effects of promising variants on gene expression, addiction characteristics, brain structure (gray and white matter) and addiction behaviors in addiction animal models (chronic administration and self-administration) were assessed. In addition, we assessed the genetic correlation among the three SDs using LD score regression. We identified and replicated three novel loci that were associated with the common risk of heroin, methamphetamine addiction and alcoholism: *ANKS1B* rs2133896 (P_meta_=3.60×10^−9^), *AGBL4* rs147247472 (P_meta_=3.40×10^−12^) and *CTNNA2* rs10196867 (P_meta_=4.73×10^−9^). Rs2133896 in *ANKS1B* was associated with *ANKS1B* gene expression and had effects on gray matter of the left calcarine and white matter of the right superior longitudinal fasciculus in heroin dependence. Over-expression of *anks1b* gene in the ventral tegmental area decreased addiction vulnerability for heroin and methamphetamine in self-administration rat models. Our findings could shed light on the root cause for substance dependence and will be helpful for the development of cost-effective prevention strategies for general addiction disorders.

## Introduction

Substance dependence (SD), which includes prescribed or illicit drug use, is a complex disease that is affected by genetics, environmental factors and pharmacological effects. The prevalence of SD to a great extent is affected by the availability of addictive substances and the national culture. Currently, nicotine, alcohol, cannabis, opioid and amphetamine-type stimulants are the most commonly used addictive substances worldwide[1]. However, the transition to different types of SD is common as several new types of addictive substances emerge. The occurrence of cross-tolerance and cross-dependence between different types of SD is high[2]. Hence, cost-effective preventive strategies for the diverse types of SD are desired.

Different addictive substances have several active mechanisms, however they converge on the reward pathways and have similar changes for cellular and molecular adaptations[3]. Individuals, who are susceptible to drug exposure and become dependent have several common characteristics, such as high impulsivity/novelty seeking and behavioral disinhibition[4]. Genetics contribute to about 40%–60% of SD vulnerability [5]. Among that, generalized genetic vulnerability may attribute to at least 20% etiology for the different types of SD [6] and may also contribute to the high comorbidity for poly-substance abuse [7,8]. Hence genetics is an important way to assess common neural and molecular mechanisms for addictive disorders. Several studies support the notion of shared genetic effects across different types of SD[9]. Through converging the varying results from previous studies, Uhl et al. found that addiction-related genes are mostly related to cell adhesion and memory processes[10]; Li et al. found that genetic variants for several genes, i.e. aldehyde dehydrogenases, *GABRA2*, and *ANKK1*, were strongly associated with dependence to various substances[11]. By performing gene network analysis on previously published genetic findings, Reyes-Gibby et al. identified *ERK1/2* to be strongly linked to smoking, alcohol, and opioid addiction[12]. However, direct evidence for these common genetic addiction factors is still lacking.

In this study, we aimed to assess the common genetic risk factors for alcoholism, methamphetamine and heroin dependence, which are among the most commonly used addictive substances. We then tried to decipher the mechanism of this genetic association.

We first present a two-stage and case-control genome-wide association analysis (GWAS) in a combined cohort for alcoholism, heroin and methamphetamine dependence. Then we sequentially examined the genetic effects of any promising cross-addiction variants on gene expression, addiction characteristics and brain images of SD patients, and addictive behaviors in drug dependence models. In addition, we assessed genetic correlation among the three SDs using LD score regression.

## Methods

### GWAS discovery cohorts

All study subjects were of Chinese Han ethnicity and were older than 16 years of age and able to understand the contents of the questionnaire. This study was approved by the Peking University Institutional Review Board and was performed in accordance with the relevant guidelines and regulations. All individuals signed a written informed consent form and were paid for their participation.

We recruited 1,026 heroin abusers (737 males, 289 females), 1,749 methamphetamine (MA) abusers (1,255 males, 494 females) and 521 male alcoholic inpatients from drug addiction treatment centers and psychiatric hospitals in China. 2,859 healthy controls (HCs) were recruited from local communities through advertisements and community centers. Heroin dependence and alcoholism was defined using the DSM-IV criteria. MA was defined using the DSM-V criteria. Basic information and substance use characteristics (e.g., age onset, dosage, frequency) were recorded. In addition, patients with alcohol dependence were required to complete the Michigan Alcoholism Screening Test (MAST). Study patients had no history of poly-substance abuse, except for nicotine dependence. All study subjects had no history of other major psychiatric disorders and medical conditions (e.g., cardiovascular, endocrinological, oncological, or autoimmune diseases) based on self-reports. Additional details regarding subject recruitment are described in the supplemental methods.

### Genetic replication cohorts

The replication sample included 1,045 male heroin abusers, 763 male methamphetamine and 146 male AD inpatients and 1,904 male HCs. The inclusion criteria (except for updating the DSM-V criteria for heroin dependence and alcoholics) and exclusion criteria were consistent with the discovery (detailed in the supplemental materials).

### Genotype and imputation

Genomic DNA was extracted using 5 mls of peripheral blood from all study participants and were used for genotyping. Genotyping for the discovery cohort was performed using the Illumina Global Screening Array-24 v1.0 BeadChip (Illumina, Inc., San Diego, CA, USA). Quality control for the SNP and individuals are described in the supplemental methods. After quality control, the final dataset included 383,065 SNPs for discovery analysis. Genotype imputation in the discovery sample were performed using the pre-phasing/imputation stepwise approach in IMPUTE2 and SHAPEIT. The imputation reference set consisted of 2,186 phased haplotypes from the full 1000 Genomes Project Integrated Phase 1 Release (March 2012). Imputed SNPs with info < 0.6 or SNPs with minor allele frequency < 0.01 were removed. Significant SNPs from the discovery stage were genotyped in the replication cohort using the Agena MassArray Analyzer 4.0 (Agena Bioscience, Inc., San Diego, CA, USA).

### Statistical analysis

Genome-wide association analysis for the discovery stage was performed using SNPTEST (-frequentist 1, -method score) (https://mathgen.stats.ox.ac.uk/genetics_software/snptest/snptest.html) for the data after imputation. Gender, age and top 10 principle components (PCs) from principle component analysis (PCA) were used as covariates. Genome-wide significance was considered as *P* < 5 × 10^−8^. Manhattan and Q-Q plots were generated using the R package, qqman. Regional plots were generated using LocusZoom. Association analysis for the replication stage cohort was performed with PLINK 1.9 using an additive model in logistic regression, with age as the covariate. The sample size weighted meta-analysis in METAL was used for analysis of the two stages. The genetic correlation among the three SDs were performed using LD score regression from the GWAS summary data as input[13]. LD scores for East Asians were downloaded from the LDSC website (https://data.broadinstitute.org/alkesgroup/LDSCORE/eas_ldscores.tar.bz2). For details of the bioinformatics analysis, please refer to the supplemental materials.

### Genetic effects on addiction characteristics

To investigate the association of the significant locus in chr12 *ANKS1B* with addiction characteristics (MAST/age onset/dosage in alcohol; age onset/frequency/dosage in methamphetamine and heroin group separately), we calculated a gene risk score (GRS) for all SNPs with P<10^−6^ and LD r^2^ > 0.75 with the lead SNP from the combined GWAS in this locus by using the --score command in PLINK. This used the BETA value of the combined GWAS as the SNP score. The GRS association with addiction characteristics were analyzed using a linear regression model, adjusted for age, gender and 10 PCs for the GWAS association analysis. Since our patients were concurrent nicotine users, we also assessed the effects of the significant SNPs on smoking phenotypes (cigs per day/ever smoking/age onset) by referring to GWAS summary data derived from 1.2 million individuals[14].

### MRI acquisition and analysis

Magnetic resonance imaging (MRI) scans and analysis were performed for a subset of patients in the heroin discovery cohort, which consisted of 65 male heroin patients and 69 male healthy controls. They all had education levels that were higher than primary school. Additional exclusion criteria for imaging included left-handedness and contraindications for MRI.

MRI data were acquired using a GE Signa Twin speed MRI 1.5T scanner (General Electric Medical System, Milwaukee, WI, USA) and a standard 8-channel head coil at 2011. A voxel-based morphometry (VBM) analysis was used to compare the whole-brain gray matter volume of all subjects using the VBM8 software package (http://dbm.neuro.uni-jena.de/vbm/). Diffusion image preprocessing and analysis were performed using the FMRIB’s Diffusion Toolbox (FDT) (FSL 4.1.4; www.fmrib.ox.ac.uk/fsl). (Detailed in supplemental methods)

### Drug addiction model

We assessed the effects of the identified target genes on addictive behaviours using chronic drug administration and self-administration models. Our estimates of the number of animals that was needed for the behavioral tests were based on previous experience in our laboratory and previous studies. All animal procedures were performed in accordance with the National Institutes of Health Guidelines for the Care and Use of Laboratory Animals and approved by the Local Animal Care and Use Committee.

For chronic drug administration experiments, rats (n=6 per group) were injected daily with methamphetamine or heroin for 16 consecutive days to assess *anks1b* expression changes after drug administration. Timelines are presented in Figure 3A. After 16 days of drug injection, rats were decapitated, and their brains were harvested promptly for western blot assays. Details of procedures are found in the supplemental methods.

To determine whether over-expression of *anks1b* could regulate addiction vulnerability, we injected a recombinant adeno-associated virus (rAAV) vector over-expressing the *anks1b* gene or a control virus into the ventral tegmental area (VTA) of rats (n=8-10 per group). VTA is the most critical brain area for addiction[15]. After virus infection, rats were trained to self-administer methamphetamine or heroin (12 training days). The dose response test (5 injection dose sessions) was then administered after training. Detailed procedures for self-administration training were described in our previous studies[16,17] as well as in Figure 3 and the supplemental methods.

## Results

### Sample characteristics

The three cohorts for substance dependence (521 alcohol, 1,749 methamphetamine, 1,026 heroin) and 2,859 healthy controls were used for GWAS. The independent replication cohort included 1,954 SD patients (1,045 heroin abusers/763 MA abusers/ 146 alcoholics) and 1,904 controls. The demographics and addiction characteristics (e.g., age onset, dosage, frequency and smoking) of the discovery and replication cohorts are presented in Table 1.

**Table 1.** Demographic and addiction characteristics of the discovery and validation cohorts.

|  | Discovery cohort |  |  |  | Replication cohort |  |  |  |
| --- | --- | --- | --- | --- | --- | --- | --- | --- |
|  | MA | Heroin | Alcohol | Control | MA | Heroin | Alcohol | Control |
| N | 1749 | 1026 | 521 | 2859 | 763 | 1045 | 146 | 1904 |
| Gender (male/female) | 1255/494 | 737/289 | 521/0 | 1851/1008 | 763/0 | 1045/0 | 146/0 | 1904/0 |
| Age (year) | 30.51 ± 7.40 | 35.6 ± 6.95 | 45.17 ± 9.27 | 34.17 ± 6.41 | 36.36 ± 8.52 | 40.06 ± 7.93 | 41.79 ± 8.90 | 33.71 ± 11.80 |
| Education level (year) | 9.34 ± 2.35 | 8.31 ± 2.69 | 10.37 ± 2.80 | N.D. | 8.25 ± 2.54 | 7.66 ± 2.33 | 11.08 ± 3.20 | N.D. |
| Smoker (%) | 88.1% | 97.1% | 94.9% | N.D. | 99.7% | 99.9% | 88.4% | N.D. |
| Onset age of addiction | 24.13 ± 7.00 | 23.13 ± 6.49 | 22.39 ± 6.77 | N.A. | 28.82 ± 9.00 | 26.36 ± 8.00 | 17.43 ± 4.29 | N.A. |
| Dosage (g/per-time for heroin and MA, standard drinks for alcohol<sup>a</sup>) | 0.65 ± 0.48 | 0.66 ± 0.60 | 11.78 ± 10.76 | N.A. | 0.53 ± 0.62 | 0.30 ± 0.19 | 17.14 ± 10.66 | N.A. |
| Frequency (times per day) | 1.51 ± 1.71 | 3.67 ± 2.36 | N.D. | N.A. | 2.03 ± 1.53 | 2.76 ± 1.49 | N.D. | N.A. |
| MAST<sup>b</sup> | N.A | N.A. | 10.22 ± 5.14 | N.A. | N.A. | N.A. | 28.53 ± 10.16 | N.A |

<sup>a</sup> Standard drink was defined as 10 grams of alcohol (equivalent to 12.5 mL of pure alcohol).
<sup>b</sup> Michigan Alcoholism Screening Test (MAST) was measured for the severity of alcoholism.
N.A.: not applicable; N.D.: not data.

### GWAS for combined SDs vs controls and replication cohort

For the combined SD vs controls comparison, we found four genome-wide significant loci, lead SNPs were *ANKS1B* rs2133896 (P=4.09×10^−8^), *AGBL4* rs147247472 (P=4.30×10^−8^), *CTNNA2* rs10196867 (P=4.67×10^−8^) and *ADH1B* rs1229984 (P=6.45×10^−10^) (Table 2). The Manhattan and Q-Q plots for the GWAS results are shown in Figure 1A and 1B respectively. The regional plots for each significant locus are shown in Figure 1C-1F.

**Table 2.** GWAS analysis and validation results.

| SNP | Chr | Minor allele | MAF<sup>a</sup> | Gene | Addiction | Discovery |  |  | Replication |  |  | Meta |  |  |
| --- | --- | --- | --- | --- | --- | --- | --- | --- | --- | --- | --- | --- | --- | --- |
|  |  |  |  |  |  | BETA | SE | P | BETA | SE | P | BETA | SE | P |
| rs2133896 | 12 | T | 0.2215 | ANKS1B | All | 0.102 | 0.019 | 4.09×10<sup>-8</sup> | 0.129 | 0.058 | 0.026 | 0.104 | 0.018 | 3.60×10<sup>-9</sup> |
|  |  |  |  |  | Alcohol | 0.233 | 0.116 | 0.044 | -0.128 | 0.161 | 0.426 | 0.110 | 0.094 | 0.242 |
|  |  |  |  |  | MA | 0.093 | 0.020 | 3.44×10<sup>-6</sup> | 0.204 | 0.074 | 5.79×10<sup>-3</sup> | 0.100 | 0.019 | 1.97×10<sup>-7</sup> |
|  |  |  |  |  | Heroin | 0.080 | 0.019 | 4.30×10<sup>-5</sup> | 0.092 | 0.071 | 0.192 | 0.081 | 0.019 | 1.78×10<sup>-5</sup> |
| rs147247472 | 1 | A | 0.0174 | AGBL4 | All | 0.286 | 0.052 | 4.30×10<sup>-8</sup> | 1.051 | 0.175 | 2.03×10<sup>-9</sup> | 0.348 | 0.050 | 3.40×10<sup>-12</sup> |
|  |  |  |  |  | Alcohol | 0.742 | 0.360 | 0.040 | N.D.<sup>b</sup> | N.D.<sup>b</sup> | N.D.<sup>b</sup> | - | - | - |
|  |  |  |  |  | MA | 0.222 | 0.055 | 5.89×10<sup>-5</sup> | 1.038 | 0.199 | 1.72×10<sup>-7</sup> | 0.281 | 0.053 | 1.34×10<sup>-7</sup> |
|  |  |  |  |  | Heroin | 0.215 | 0.052 | 4.05×10<sup>-5</sup> | 1.078 | 0.197 | 4.66×10<sup>-8</sup> | 0.272 | 0.051 | 7.88×10<sup>-8</sup> |
| rs10196867 | 2 | C | 0.0297 | CTNNA2 | All | 0.239 | 0.044 | 4.67×10<sup>-8</sup> | 0.275 | 0.129 | 0.033 | 0.243 | 0.042 | 4.73×10<sup>-9</sup> |
|  |  |  |  |  | Alcohol | 0.610 | 0.272 | 0.025 | N.D.<sup>b</sup> | N.D.<sup>b</sup> | N.D.<sup>b</sup> | - | - | - |
|  |  |  |  |  | MA | 0.216 | 0.046 | 3.30×10<sup>-6</sup> | 0.098 | 0.171 | 0.565 | 0.208 | 0.045 | 3.50×10<sup>-6</sup> |
|  |  |  |  |  | Heroin | 0.181 | 0.045 | 4.83×10<sup>-5</sup> | 0.409 | 0.147 | 5.46×10<sup>-3</sup> | 0.200 | 0.043 | 2.67×10<sup>-6</sup> |
| rs1229984 | 4 | C | 0.299 | ADH1B | All | 0.105 | 0.017 | 6.45×10<sup>-10</sup> | - | - | - | - | - | - |
|  |  |  |  |  | Alcohol | 0.677 | 0.095 | 1.03×10<sup>-12</sup> | 2.038 | 0.156 | 4.80×10<sup>-39</sup> | 1.045 | 0.081 | 5.32×10<sup>-38</sup> |
|  |  |  |  |  | MA | 0.000 | 0.019 | 0.986 | N.D.<sup>c</sup> | N.D.<sup>c</sup> | N.D.<sup>c</sup> | - | - | - |
|  |  |  |  |  | Heroin | 0.001 | 0.019 | 0.942 | N.D.<sup>c</sup> | N.D.<sup>c</sup> | N.D.<sup>c</sup> | - | - | - |

N.D.: No data. MA: methamphetamine. <sup>a</sup> Minor allele frequency (MAF) of the SNP in the discovery stage. <sup>b</sup>rs147247472 and rs10196867 had too low MAF to be tested in the small validation sample (n=146) for alcoholism, hence it was not tested. <sup>c</sup>ADH1B rs1159918 was not significant for MA and heroin in discovery stage, hence it was not tested in MA and heroin groups in the validation cohort.

**Figure 1.**
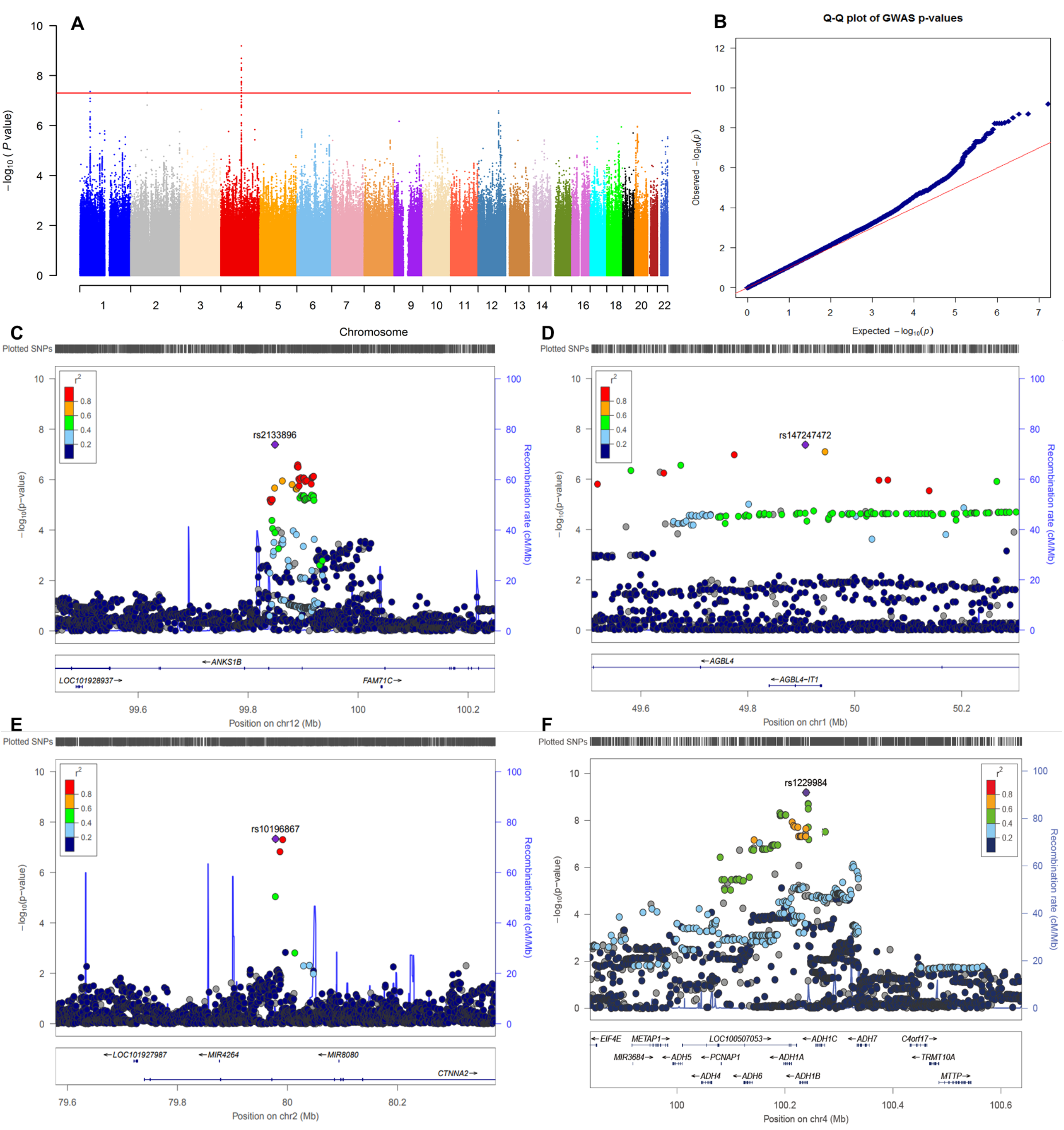
GWAS association analysis. (A) Manhattan plot for the GWAS of the combined substance dependence cohort. (B) Q-Q plot for the combined substance dependence GWAS p-values. Regional plots for the four genome-wide significant loci: *ANKS1B* rs2133896 (C), *AGBL4* rs147247472 (D), *CTNNA2* rs10196867 (E) and *ADH1A* rs1229984 (F).

We then checked the genetic association for these significant SNPs for each of the three addictions vs controls. *ANKS1B* rs2133896, *AGBL4* rs147247472, and *CTNNA2* rs10196867 were synchronously associated with alcoholism, heroin and methamphetamine dependence respectively (P<0.05, Table 2), while *ADH1B* rs1159918 was only significant for alcoholism (P=1.03×10^−12^ for alcohol, P=0.986 for MA and P=0.942 for heroin).

For the replication cohort, the association between *ANKS1B* rs2133896 and SD was validated in the combined cohort for alcohol, methamphetamine and heroin dependence (P_replication_ = 0.026, P_meta_ = 3.6×10^−9^). For *AGBL4* rs147247472 and *CTNNA2* rs10196867, their association with SD was replicated in the combined methamphetamine and heroin cohorts (P_replication_ = 3.4×10^−12^ and 4.73×10^−9^ respectively). Due to their low MAF (0.0174 and 0.0297 respectively), they were hardly tested in our relatively small alcoholism cohort (n=146) (Table 1). *ADH1B* rs1159918 was only tested in the alcoholic cohort and was replicated to be associated with alcohol dependence (P_replication_=4.8×10^−39^, P_meta_=5.32×10^−38^).

### Gene-level and gene-set level analysis of GWAS results

We wanted to understand the common genetic risks of SD at the gene and pathway level. Gene-level analysis for the combined GWAS data identified three significant genes (*ADH1A, ADH1B* and *ADH6*). The top 10 genes are shown in Table S1. Gene-set analysis found two significant pathways “Wnt signaling pathway” and “Basal cell carcinoma” that passed multiple corrections (Table S2).

### Functional analysis of the rs2133896 locus

*ANKS1B* rs2133896 was the only SNP that was associated with all the three types of SD. It was validated in the combined cohort. The sequential analysis of e-QTL, genetic-addiction characteristics, genetic-imaging and the animal model study only focused on *ANKS1B* rs2133896 (MAF=0.222, protective allele (G), risk allele (T)).

The 15-core chromatin state data from the RoadMap project for the *ANKS1B* rs2133896 locus showed that the top lead SNP was in the enhancer region (Figure S2). This denoted that the locus may have regulatory function. We checked eQTL data for this locus. The top SNP rs2133896 was not found in BRAINEAC, hence we used its LD-proxy rs10860447 (r^2^ = 0.902, calculated using --ld in PLINK) to assess the effect of *ANKS1B* rs2133896 on gene expression. rs10860447 C allele (linked with rs2133896 T allele) had higher *ANKS1B* expression in white matter and lower expression in cerebellum cortex compared to the T allele (Figure S3A, P_uncorrected_<0.05). Using GTEx, we found that *ANKS1B* was specifically expressed in brain tissue (Figure S3B).

### Association of the *ANKS1B* locus with addiction characteristics

We then assessed the effects of GRS of *ANKS1B* locus on addiction characteristics for each type of SD. There was no significant association between *ANKS1B* GPS with addiction characteristics. We found only a nominal genetic association between *ANKS1B* GPS heroin use frequency (BETA = 5.78, P_uncorrected_ = 0.035) (Table S3). In addition, we did not find a significant association for *ANKS1B* locus SNPs with smoking phenotypes from a large GWAS [14].

### Association of *ANKS1B* rs2133896 with imaging characteristics

We assessed the effect of rs2133896 on the gray matter and found a significant interaction effect of *ANKS1B* rs2133896 (TT genotype and G carriers) × group (heroin and healthy) on the gray matter volume of the left calcarine (CAL.L) (x=-27 mm, y=-64 mm, z=7 mm, voxel size=491; d=0.075, P_AlphaSim-corrected_<0.001, 95% CI=0.032-0.136, Figure 2). *Post hoc* test results showed that the gray matter volume of heroin patients who carried the risk TT genotype was significantly decreased in CAL.L compared to G allele carriers of heroin patients. Although there was a lack of statistically significance, the CAL.L gray matter volume of healthy controls who carried the risk TT genotype showed an increased trend compared to healthy G allele carriers.

**Figure 2.**
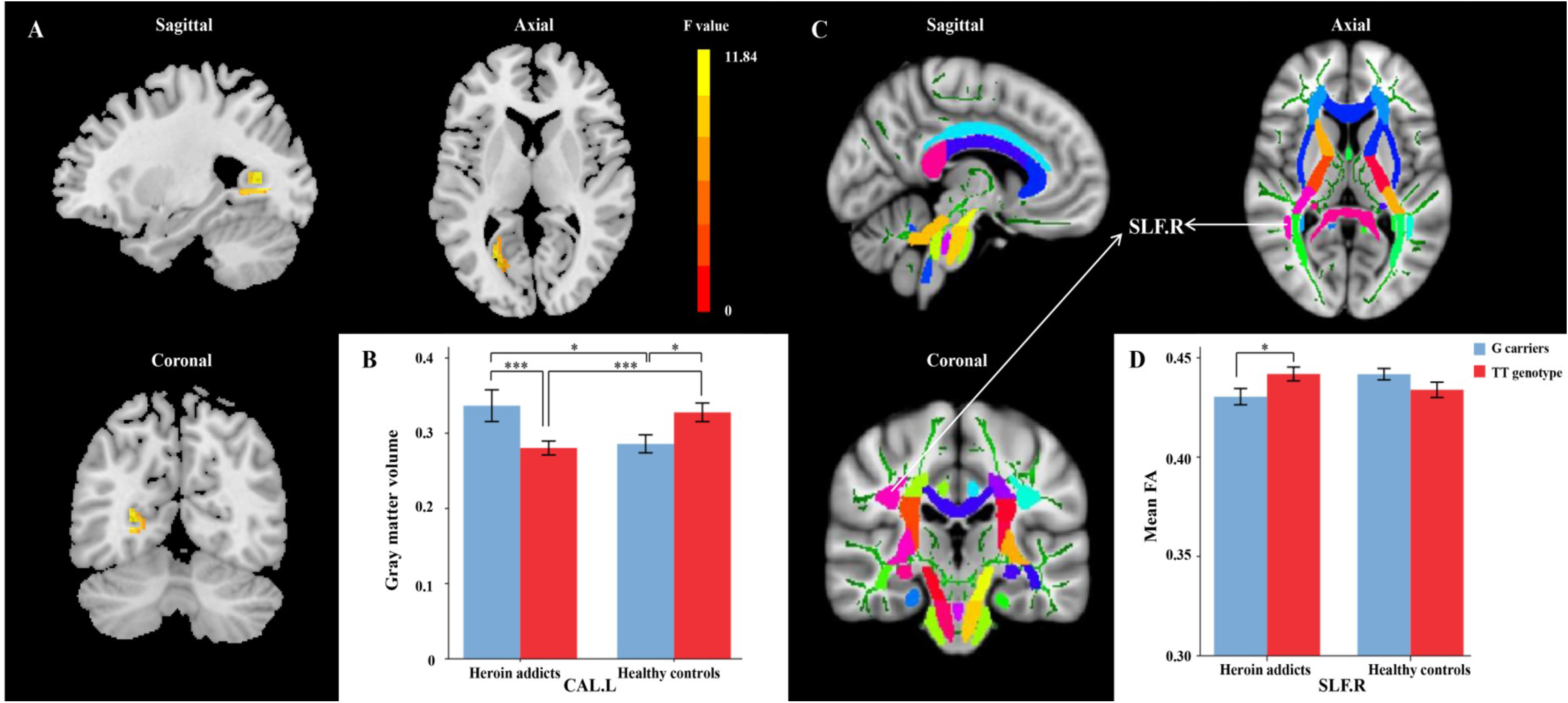
Interaction effects of ANKS1B rs2133896 by heroin on brain imaging. (A) Significant interaction effect of *ANKS1B* rs2133896 (TT genotype and G carriers) × group (heroin and healthy) on the gray matter volume of the left calcarine (CAL.L) (B) *Post hoc* test results for CAL.L. (C) After Bonferroni correction, significant interaction effect of genotype × group was found in the right superior longitudinal fasciculus (SLF). (D) *Post hoc* test results for right SLF.

We then assessed the effect of rs2133896 on twenty major tracts of white matter. After Bonferroni correction, significant interaction effect of *ANKS1B* rs2133896 × group was found in the right superior longitudinal fasciculus (SLF) (F=10.43, P=0.002). *Post hoc* test results showed that, compared to G allele carriers, the mean FA of the right SLF was significantly increased in risk TT genotype carriers of heroin abusers (Figure 2, Table S4).

### Role of *anks1b* gene in drug addiction models

In chronic drug administration experiments, we found that *anks1b* expression in VTA was significantly decreased in both methamphetamine (t_8_=3.411, P=0.009) and heroin administered rats (t_10_=3.085, P=0.02) compared to saline treated rats (Figure 3A and B). This suggests that addictive drugs could suppress *anks1b* gene expression. Over-expression of *anks1b* may reverse addictive behavior.

**Figure 3.**
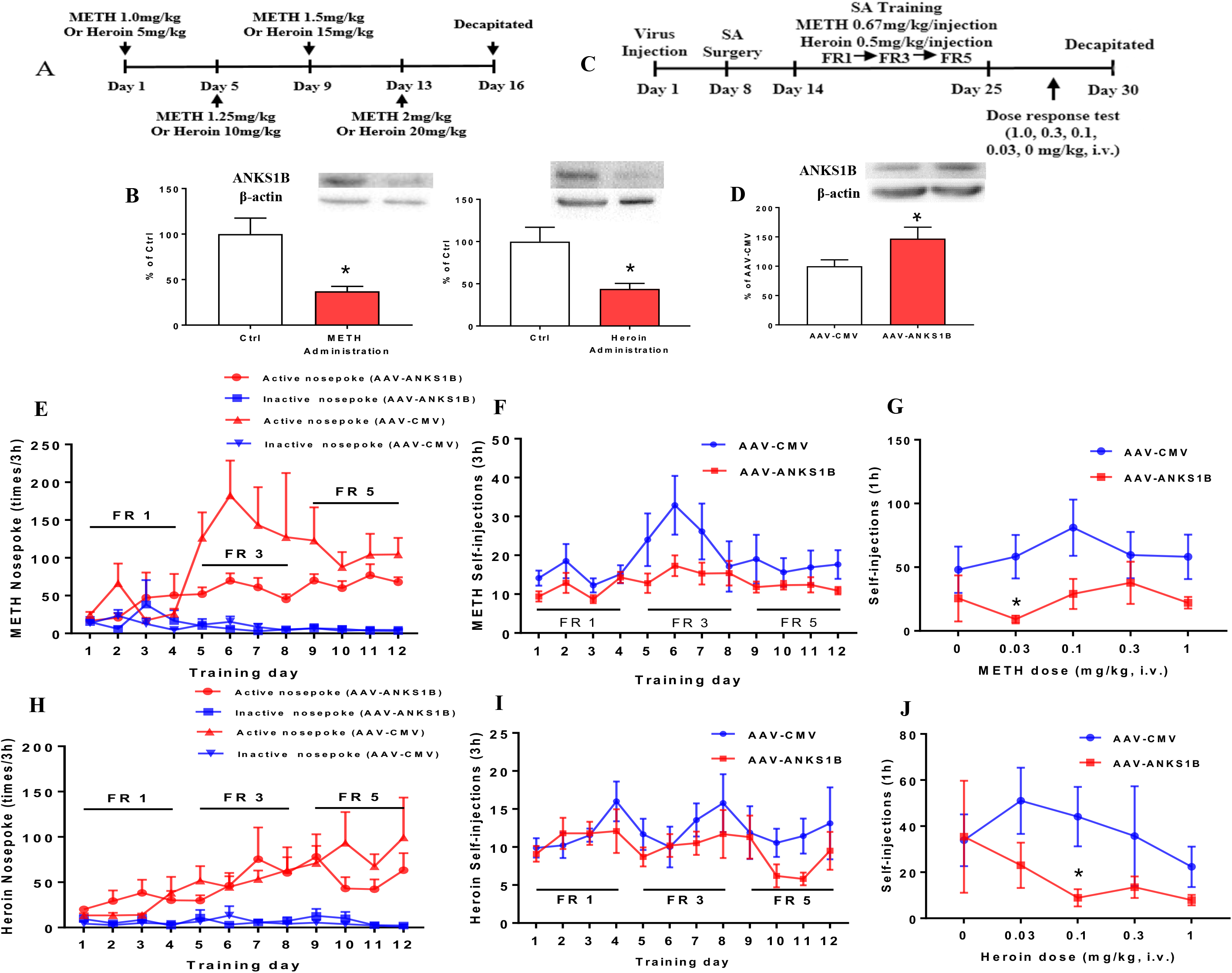
Role of *anks1b* in drug addiction animal models. (A) Timeline of chronic drug administration procedure. (B) Western blot analysis of *anks1b* in VTA after 16 days of drug treatment. (C) Timeline of the drug self-administration procedure. (D) Western blot analysis of *anks1b* in VTA from rats injected with AAV-ANKS1B or AAV-CMV. (E) Number of responses (mean ± SEM) on the active and inactive nose-poke devices during methamphetamine self-administration training sessions. (F) Number of self-injections during methamphetamine self-administration training. (G) Dose response test for 5 injection-dose methamphetamine sessions. (H) Number of responses on the active and inactive nose-poke devices during heroin self-administration training sessions. (I) Number of self-injections during heroin self-administration training. (J) Dose response test for 5 injection-dose heroin sessions. *P<0.05 compared to the control.

During self-administration training, we found no significant difference between *anks1b* over-expressing rats and control rats (for both methamphetamine and heroin training) in terms of the total number of activated nose pokes and self-injections. (Figure 3E, F, H and I). For the dose-response test, compared to the controls, *anks1b* over-expressed rats had a significantly lower number of injections for both methamphetamine (0.03mg/kg, t _(1,16)_=3.147, P=0.006) and heroin (0.1mg/kg, t _(1,17)_=2.750, P=0.014) (Figure 3G and J). Repeated measures ANOVA (2 groups × 5 different doses) revealed that the main effect was group (*anks1b* over-expressed vs control) on response rates: for methamphetamine group (F _(1,16)_=13.965, P <0.0001) and for heroin group (F _(1,17)_=5.530, P=0.021). This suggests that over-expression of *anks1b* could decrease the addiction vulnerability in self-administrated rats.

### Genetic correlation between the three SDs

To further understand the genetic association between the three SDs, we analyzed the genetic correlation using LD score regression analysis. LD score regression analysis for the GWAS summary of the three SDs showed that the SNP heritability for alcohol, heroin and MA dependence were h^2^=0.169 (SE=0.1378), h^2^=0.2211 (SE=0.099) and h^2^=0.1773 (SE=0.1361) respectively. Genetic correlation analysis showed that heroin dependence had a high genetic correlation with MA (r_g_ = 0.603), but the significance was only a trend (P=0.077). Genetic correlations between alcohol dependence and heroin dependence, alcohol dependence and MA dependence were not significant.

## Discussion

This is the first GWAS analyses performed to identify critical genetic etiologies associated with different types of SDs. This was performed using a combined cohort of alcohol, heroin and methamphetamine dependent users. We found three specific loci (peaked in *ANKS1B* rs2133896, *AGBL4* rs147247472, and *CTNNA2* rs10196867) that were synchronously associated with alcoholism, heroin and methamphetamine dependence. The genetic association between *ANKS1B* rs2133896 and SD was validated in combined replication cohorts. In addition, we found that rs2133896 affected *ANKS1B* gene expression, use frequency and brain imaging changes in heroin patients. Moreover, the role of *ANKS1B* gene in drug addiction was underscored in rat model experiments.

Our findings were partly consistent with the converging results of addiction genetic studies, which also showed that *ANKS1B* variants contributed to risk to multiple types of SDs[10]. Gene-set convergence results suggested that the “Wnt signaling pathway”, which is involved in the occurrence and development of psychiatry disorders[18], could play a critical role in the common risk of SD. Of note is the association between *ANKS1B* variants with alcoholism, which did not replicate well. Our LD score regression results suggested the genetic relation between MA dependence and heroin dependence may be closer compared to that of alcoholism. This may be due to alcoholism having specific metabolic enzymes and relatively weaker effects on reinforcing properties compared to illegal drug dependence [19,20]. The key genetic components in alcohol metabolizing enzyme genes (e.g., *ADH* and *ALDH* variants) were reliably and specifically correlated with alcoholism[8,21]. Hence, the cross-addiction related genetic factors may have relatively uncertain association with alcoholism and needs further genetic studies using larger sample cohorts and mechanistic studies to substantiate their association. Additionally, most of our patients were also smokers, which made smoking a critical confusing factor. The effects of promising variants on smoking were difficult to discriminate because we lacked data regarding smoking in healthy controls. In previous GWAS on nicotine use, we found no evidence of SNPs related with smoking. In addition, we did not find significant association between these promising SNPs with smoking phenotypes using a large GWAS dataset. Hence, we believe that the confusing effects of smoking would be limited.

*ANKS1B* gene (aliases: EB-1, AIDA-1) is predominantly expressed in the brain and encodes an activity-dependence effector of post-synaptic signaling[22,23]. It may act as a synapse-to-nucleus messenger in the regulation of synaptic plasticity and control of protein biosynthetic capacity (GO Source: InterPro)[24]. *ANKS1B* variants have also been shown to modulate monoamine metabolite levels in the cerebrospinal fluid [25] that play a role in psychiatry disorders. Homozygous *anks1b* KO genotype is partially lethal and exhibits locomotor hyperactivity and increased stereotypy[26]. In previous studies, *ANKS1B* variants were associated with common genetic risks that shared effects on five major psychiatric disorders[27], and were among the top hits in several antipsychotic drug response GWAS[28–30]. Our findings regarding *ANKS1B* variants associating with different types of SD extend its important role in the genetic etiology of psychiatry disorders. Genetic-phenotypes results further suggest the effects of *ANKS1B* rs2133896 enhancer variant, on drug dependence, and may be mediated by affecting the brain structure in CAL and SLF. The visual cortical, of which the center is CAL, is the main sensory association that is necessary for reward processing[31] and cue-induced drug cravings[32]. SLF is the critical association fiber tract that links the frontal, occipital, parietal and temporal lobes[33]. Hence, we speculated that patients who carry the rs2133896 risk genotype may facilitate the transition of drug-related sensory stimulus though abnormal white matter connectivity in SLF to aggravate addiction development.

In addition, we identified two significant loci with low MAF, i.e., *AGBL4* rs147247472 and *CTNNA2* rs10196867. *AGBL4* is an ATP/GTP binding protein like 4 gene and is expressed in the brain (supplemental Figure S4A). The age-related methylation changes in *AGBL4* are associated with cognitive function[34]. Whereas rs147247472 located in the *AGBL4* intron region has been rarely reported. Other variants of *AGBL4* have been reported in several SD GWAS [35,36]. *CTNNA2* encodes a cell-adhesion protein-catenin alpha 2 and is strongly expressed in the central nervous system, predominately in the prefrontal cortex (Supplementary Figure S4B). *CTNNA2* is engaged in regulating synaptic plasticity[37], brain morphogenesis and behavior performance[38]. *CTNNA2* variants achieved genome-wide significance for attention deficit hyperactivity disorder [39] and excitement-seeking personality[40]. *CTNNA2* variants have been constantly identified in addiction GWAS[41], and hence suggests the important role of *CTNNA2* variants in addiction-related behaviors. Given the low MAF of the loci and our limited sample size, additional replication and biological mechanistic studies are necessary for bridging the association between these novel variants and addiction.

There were two limitations to our study. First, the dynamic interaction between genetic, environmental factors and pharmacologic effects was complex, hence the definition of healthy controls and patients with specific addictions could be time-specific. Second, our findings need to be replicated using other types of SD and additional ethnic groups.

In summary, our findings revealed several novel genome-wide significant SNPs and genes associated with common susceptibility and effects on phenotypic changes for alcoholism, heroin and MA dependence. These findings shed light to the root cause and categorical distinctions of different SD and will help develop early prevention measures for all SD. We would like to evaluate the interaction of common and unique genetic effects on different types of SD in our future studies.

## Contributions

J.S. Y.S. and S-H.C. designed the study and obtained financial support; Y.S., F.W., W.-H.Y., H.-Q.S., Z.-J.N. and X.-W.Z. conducted cohort recruitment, collected biological samples and phenotypic data. J-Q.L performed the genotype microarray experiments. S-H.C and Y.S. performed genetic data processing, statistical and bioinformatics analysis. Z.L. performed the imaging. L.-B.Z., Y.-B.Z. and Y.C performed the animal and in vitro experiments. Y.S and S.-H.C. drafted the manuscript. J.S and L.L. supervised the experiments and data analysis. All authors critically reviewed the manuscript and approved the final version.

## Acknowledgements

This work was supported by grants from the National Basic Research Program of China (2015CB553503), the National Natural Science Foundation of China (U180220091, 81821092, 81601165), the National Key Research and Development Program of China (2017YFC0803608, 2017YFC0803609, 2016YFC0800908), Beijing Municipal Science & Technology Commission (Z181100001518005 and Z161100002616006), and Youth Elite Scientists Sponsorship Program by CASR (CSTQT2017002). We are grateful to Beijing Compass Biotechnology Company for technical assistance with the microarray experiments.

