## supplementary materials for "Identification of novel risk loci with shared effects on alcoholism, heroin and methamphetamine dependence"

Running title: Common genetic risk factors for addiction

*Correspondence:

Prof. Jie Shi

National Institute on Drug Dependence, Peking University, 38 Xueyuan Road, Haidian District, Beijing 100191, China. Tel: 86-10-8280-1593, Fax: 86-10-62032624

**This file includes:**

Supplementary Methods

Supplementary Figure S1- S4

Supplementary Table S1-S4

### Supplemental methods

#### 1. Subjects

1) Heroin abusers: We recruited 1,026 heroin abusers (737 males, 289 females) from five drug addiction treatment centers in China's Guangdong and Hubei provinces (Yichang, Sanshui, Shenzhen and Huizhou) from Jan. 2011 to Mar. 2014. Using self-reported questionnaires and government records, heroin was their primary drug of abuse. Confirmatory urine tests were positive for heroin at the time of treatment entry. Individuals met the DSM-IV criteria for heroin dependence as evaluated by trained interviewers. Substance use characteristics (onset age for regular abuse, dose per time (g) and usage frequency per day within the past one-year) were recorded (Table S1). Study patients had no history of poly-substance abuse (continuous use of other opioid drugs for no more than one month and use of other kinds of addictive drugs for no more than three times within the past three years), apart from nicotine dependence. Alcoholism was also excluded by selecting patients with a score < 4 in the Michigan Alcoholism Screening Test. All study subjects had no history of other major psychiatric disorders and medical conditions (e.g., cardiovascular, endocrinological, oncological, or autoimmune diseases) based on self-reports.

2) Methamphetamine abusers: A total of 1,749 methamphetamine (MA) abusers (1,255 males, 494 females) were recruited from three addiction treatment centers in China's Guangdong province (Dongguan, Shenzhen and Sanshui) from Dec. 2016 to Mar. 2017. MA was their primary drug of abuse and urine tests confirmed MA use at time of treatment entry. Patients met the DSM-V criteria for MA. Substance use characteristics are showed in Table S1. Patients did not use other non-amphetamine-type stimulants (e.g., opioid and cocaine) for more than three times per year within the past three years, with the exception of nicotine. Alcoholism was also excluded by selecting patients with a score < 4 in the Michigan Alcoholism Screening Test. All study subjects had no history of other major psychiatric disorders and medical conditions based on self-reports.

3) Alcohol abusers: A total of 521 male alcoholic inpatients were recruited from twelve psychiatric hospitals in northern China (Beijing, Inner Mongolia Autonomous Region, Shandong, Tianjin, Jilin, Liaoning and Heilongjiang provinces) from Sept. 2009 to Nov. 2011. All enrolled patients sought treatment for alcohol dependence (AD) and were clinically ascertained as AD based on DSM-IV by experienced psychiatrists. The severity of alcoholism was measured by criterion counts based on the Chinese version of the Michigan Alcoholism Screening Test (MAST)[1]. Patients had not used any other addictive substances except for nicotine based on self-reports. All study subjects had no history of other major psychiatric disorders and medical conditions based on self-reports.

4) Healthy controls: 2,859 healthy controls (HCs) were recruited from local communities through advertisements and community centers. All HCs did not have a history of substance abuse (except nicotine), psychiatric disorders or any current major medical conditions based on

self-report questionnaires.

**2. Genetic replication cohorts**

1) Heroin and methamphetamine abusers: The replication sample included 1,045 male heroin abusers and 763 male methamphetamine abusers from Sanshui addiction treatment hospital in Guangdong province of China. They were enrolled in Mar. 2018. The inclusion criteria (except for updating the DSM-V criteria for heroin dependence) and exclusion criteria were consistent with the discovery cohort.

2) Alcoholics: 146 male AD inpatients were recruited from the Peking University Sixth Hospital and four other psychiatric hospitals from Anhui, Shangdong and Henan provinces in China in 2018. The inclusion (except for updating the DSM-V criteria for alcoholism) and execution criteria were similar to the discovery cohort.

3) Healthy controls: The 1,904 healthy controls (HCs) were recruited from the local community. The recruitment method and inclusion criteria were similar to the controls in the discovery cohort.

**3. Quality control for genetic analysis**

SNPs with call rates < 95%, minor allele frequency (MAF) < 0.01 or Hardy-Weinberg (H-W) equilibrium test $P < 1 \times 10^{-6}$ were removed. Samples from individuals lacking information on age or gender, call rate < 95%, autosomal heterozygosity > 5 s.d. away from the mean or being one of a pair of individuals with proportion identity by descent (IBD) PI_HAT ≥ 0.185 (the one with lower call rate) were removed. The final dataset included 3,296 cases (1,749 MA abusers, 1,026 heroin abusers, 521 alcohol abusers) and 2,859 controls with 383,065 SNPs.

Principal component analysis (PCA) was then performed for all samples. A total of 20,501 genotyped autosomal SNPs, which were in low linkage disequilibrium (LD) (MAF > 0.35 and $r^2 < 0.05$ for each pair of SNPs) and absent from the 5 long-range LD regions, were included in the PCA using EIGENSOFT 4.2 software. The PCA plots for the top four pairs of PCs are shown in Supplementary Figure 1. The top 10 principal components (PCs) were used as covariates for subsequent association analysis.

**4. Bioinformatics analysis**

Gene expression profiles of the significant genes were obtained from GTEx. e-QTL was examined using the UK Brain Expression Cohort data set (GSE46706). Expression plot was generated using BRAINEAC (http://peana-od.inf.um.es:8080/UKBECv12/). Gene-level and gene-set level association analysis were performed using the MAGMA software (Multi-marker Analysis of GenoMic Annotation, http://ctglab.nl/software/magma). The mapping window from SNP to gene was 5kb. KEGG pathway data (n = 221) was used for gene-set analysis. Significance levels were set as $0.05/18{,}225=2.74\times10^{-6}$ for gene and $0.05/221=2.26\times10^{-4}$ for gene-set separately. Chromatin state near the significant loci was obtained from FUMA[2] using the data from RoadMap[3].

**5. Imaging analysis**

**1) Imaging acquisition**

MRI data was acquired using a GE Signa Twin speed MRI 1.5T scanner (General Electric Medical System, Milwaukee, WI, USA) and a standard 8-channel head coil at Zhongshan Traditional Chinese Medicine Hospital. The subjects were in the supine position with their heads snugly fixed by straps and foam pads to minimize head movements. T1-weighted, sagittal three-dimensional images were acquired using a spoiled gradient recalled echo sequence that covered the entire brain (slice thickness = 1 mm, repetition time = 7.816 ms, echo time = 2.984 ms, inversion time = 450 ms, flip angle = 13°, acquisition matrix = 256 × 256, field of view = 256 × 256 mm2, number of averages = 2). In addition, T2-weighted images were acquired to exclude subjects with abnormal brain images and were verified by two experienced neuroradiologists. Diffusion-weighted images were acquired using a single-shot twice-refocused echo planar imaging sequence (56 axial slices, 2.4 mm slice thickness without gap, repetition time = 14.4 s, echo time = 85 ms, 25 non-linear directions with b = 1000 s/mm$^2$, three additional images without diffusion weighting [b = 0 s/mm$^2$], acquisition matrix = 128 × 128, field of view = 256 × 256 mm$^2$, number of averages = 2).

**2) Structural image analysis**

Voxel-based morphometry (VBM) analysis was used to compare the whole-brain gray matter volume of all subjects using the VBM8 software package (http://dbm.neuro.uni-jena.de/vbm/). Default parameters were used for pre-processing of structural images, except for estimation using 'ICBM space template-East Asian brains' and extended options using 'thorough clean up'. The images were bias-corrected, tissue classified and normalized to the Montreal Neurological Institute space using affine and non-linear transformations to compare the absolute amounts of tissue [4]. The modulated gray matter images were smoothed using a Gaussian kernel of 8 mm full width at half maximum (FWHM). We utilized the mean gray matter map (threshold=0.3) for all subjects to obtain a group-based brain mask and used it for subsequent analysis. A full factorial ANCOVA (2×2) was calculated using SPM8 with group (heroin addicts versus healthy controls) and the target SNP rs2133896 (G carriers versus TT genotype) as independent factors with age, cigarette use, and total intracranial volume included as covariates. The statistical threshold was set at $p<0.001$ using AlphaSim correction ($p<0.01$, >392 voxel cluster size), which was determined through Monte Carlo simulations by using the AlphaSim program (AFNI; http://afni.nimh.nih.gov/pub/dist/doc/manual/AlphaSim.pdf) [5]. The significant areas for the interaction of group×rs2133896 in the VBM analysis were extracted as regions of interest (ROI) using DPABI v3.0 (http://rfmri.org/dpabi).

**3) Diffusion image analysis**

Diffusion image preprocessing and analysis was performed using FMRIB's Diffusion Toolbox (FDT) (FSL 4.1.4; www.fmrib.ox.ac.uk/fsl). First, we corrected head movements and eddy currents. Brain masks were then created using the BET procedure on B0 (no diffusion weighting) images. Last, DTIfit was used to fit the diffusion tensor model. The output of DTIfit produced voxel-wise maps of the fractional anisotropy (FA), mean diffusivity (MD), axial diffusivity (DA), and radial diffusivity (DR). Next, we adopted the atlas-based segmentation strategy. We registered each subject's FA and MD maps to the digital white matter atlas JHU ICBM-DTI-81 (http://cmrm.med.jhmi.edu/), a probabilistic atlas generated by mapping DTI data of 134 subjects to a template image. As shown in Fig. 3, the JHU-WM atlas was overlaid on the WM skeleton of each subject in the CBM-DTI-81 space, such that each skeleton voxel could be categorized into one of the 20 major tracts. The mean diffusion metrics (i.e., FA and MD) of the twenty tracts were calculated for each subject. For the diffusion alterations in the atlas-based tract regions of interest (ROIs), we performed a MANCOVA to compare the DTI indices differences between the genotype groups, rs2133896, and interaction between the two (age, and cigarette use corrected). Bonferroni correction was used to adjust for possible spurious findings due to multiple testing ($p<0.05/20$). Subsequent post hoc comparisons were performed using the Tukey's Test.

**6. Animal model experiments**

**1) Animals**

Male Sprague Dawley rats (240-260 g) were purchased from Beijing Vital River Laboratories. The rats were maintained on a reverse 12 h/12 h light/dark cycle with controlled temperature (23 ± 2ºC) and humidity (50 ± 5%) and housed in groups of five with food and water freely available. Rats were randomly allocated into treatment condition groups.

**2) Chronic drug administration**

Methamphetamine or heroin was dissolved in 0.9% saline and injected intraperitoneally (i.p.) (n=6 per group). The rats were daily injected with methamphetamine or heroin in their cages for 16 consecutive days. For the methamphetamine administration schedule, rats were injected with different doses of methamphetamine sequentially (1.0 mg/kg, 1.25 mg/kg, 1.5 mg/kg and 2 mg/kg, respectively), with each dose injected twice per day (9:00am and 18:00pm) for 4 days. For the heroin administration schedule, rats were injected with different doses of heroin (5 mg/kg, 10 mg/kg, 15 mg/kg and 20 mg/kg, respectively), with each dose injected twice per day (9:00am and 18:00pm) for 4 days sequentially. Rats were euthanized after the last day of injection (Timelines are presented in Figure 4A).

**3) Recombinant adeno-associated virus injection**

rAAV2/9-CMV-ANKS1B-flag-pA (5.00 x $10^{12}$vg/mL) and rAAV2/9-CMV-flag-WPRE-pA (5.00 x $10^{12}$vg/mL) were generated by BrainVTA.Co. All viral vectors were stored in aliquots at -80°C until use. Approximately 1 μL of virus was injected into the ventral tegmental area (VTA) (−5.2 mm from bregma, +1.8 mm lateral to the midline (10° angles), and −7.5 mm from the skull surface) per side. The syringe was left ten minutes after the end of the injection to allow for virus diffusion. After virus administration, rats were returned to their cages for postsurgical recovery for 1 week before intravenous drug self-administration. Rats without proper virus targeting were excluded from the study. The virus injections significantly increased the expression of anks1b in VTA (t12=-2.244, P=0.045).

**4) Intravenous drug self-administration, training and testing in rats**

Rats were anaesthetized (weighing 300–320 g when surgery began) with Isoflurane. For self-administration methamphetamine training, there were 10 rats in the *anks1b* over expression group and 8 rats in the control group. For self-administration heroine training, there were 10 rats in the anks1b over-expression group and 9 rats in the control group. Catheters were then inserted into the right jugular vein with the tip terminating at the opening of the right atrium as previously described [6]. Rats were allowed to recovery for 3-5 days after surgery.

The chambers (AniLab Software & Instruments) were equipped with two nose-poke operandi (AniLab Software & Instruments) located 5 cm above the floor of the chambers. Nose pokes in one (active) operand led to drug infusions that were accompanied by a 5-s tone-light cue. Nose pokes in the other (inactive) operand were also recorded but had no consequence. Rats were trained to self-administer methamphetamine (0.67 mg/kg per infusion) or heroin (0.5 mg/kg per infusion) during the three 1-h daily sessions (separated by 5min) over 12 days. The sessions began at the onset of the dark cycle. We used a reinforcement schedule of fixed ratio 1 (FR1, each activate nosepoke delivered with one injection of drug) for the first 4 days, FR3 (three activate nosepokes delivered with one injection of drug) for the middle 5-8 days, and FR5 (five activate nosepokes delivered with one injection of drug) for the last 4 days with a 40-s timeout period after each infusion. Each session began with the house light illuminated and remained on for the entire session. We limited the number of drug infusions to 15 per hour to prevent overdose.

After self-administration training, rats were subjected to a between-session dose-response test with each of 5 injection doses (0, 0.03, 0.1, 0.3, 1 mg/kg per infusion). Each dose was tested for 1 hour and the number of injections was recorded during test sessions.

**5) Western blot assays**

Western blot assays were performed as previously described [7]. After decapitation, rat brains were promptly removed and then frozen at -60 °C N-hexane and then transferred to an -80 °C freezer. Total protein from bilateral tissue punches (12 gauge) of the VTA were extracted in ice-cold RIPA lysis buffer (APPLYGEN Biotechnology, Beijing, China; 50mM Tris-HCl, 150mM NaCl, 1% NP-40, 0.1% SDS, pH 7.4) supplemented with protease inhibitor cocktail. After homogenizing using an electrical disperser (Wiggenhauser, Sdn Bhd), the tissue lysates were centrifuged at 12,000r for 10 min at 4 °C and protein concentrations were determined from the supernatants. 5X loading buffer (16% glycerol, 20%-mercaptoethanol, 2% SDS and 0.05% bromophenol blue) was then added to each sample (4:1, sample: loading buffer) before boiling for 5min. The samples were then cooled and subjected to SDS–polyacrylamide gel electrophoresis (10% acrylamide/0.27% N, N0-methylenebisacryalamide resolving gel) for 40 min at 80V in stacking gel and 1 h at 130V in resolving gel. For each electrophoresis run, increasing amounts of protein were pooled from the brain regions being tested to produce a standard curve. Proteins were transferred to Immobilon-P membranes (Millipore) at 0.25A for 1.5 h. The membranes were then washed with TBST (Tris-Buffered Saline plus 0.05% Tween-20, pH 7.4) and then placed in blocking buffer (5% bovine serum albumin (BSA in TBST) overnight at 4 °C.

The membranes were then incubated for 1 h at room temperature on an orbital shaker with anti-ANKS1B (PA5-45780, 1:1,000; Thermo Fisher) and β-actin (TA-09, 1:2,000; ZSGB-BIO) in TBST plus 5% BSA and 0.05% sodium azide. After three 5-min washes in TBST buffer, the blots were incubated for 45 min at room temperature on a shaker with horseradish peroxidase conjugated secondary antibody (goat anti-rabbit IgG and goat anti-mouse IgG; ZB-2301 and ZB-2305; ZSGB-BIO) diluted 1:2,000 in blocking buffer. The blots were then washed three times for 5min each in TBST and incubated with Super Signal Enhanced chemiluminescence substrate (Detection Reagents 1 and 2, 1:1 ratio, Pierce Biotechnology) for 1 min at room temperature. The membranes were then removed and wrapped in plastic wrap (no bubbles between blot and wrap) and protein bands were detected using the ChemiDoc MP (Bio-Rad). Protein band intensities were quantified using the Quantity One software (version 4.4.0, Bio-Rad). Band intensities from each test sample were compared to the band intensities from the standard curves. The protein amounts of interest from each sample were interpolated from the standard curve. Test samples were within the linear range of detection.

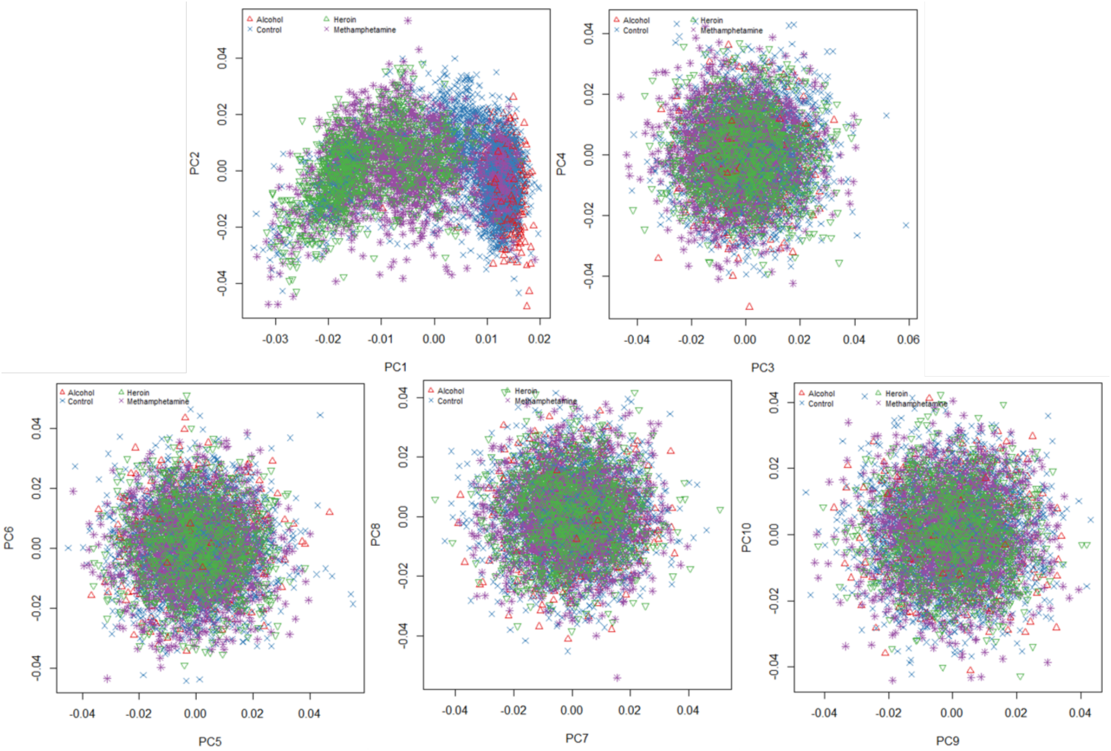


**Supplementary Figure** 1 Principle component analysis for all substance dependence cohorts and controls using smartpca (EIGENSTRAT). The first ten PCs were plotted.

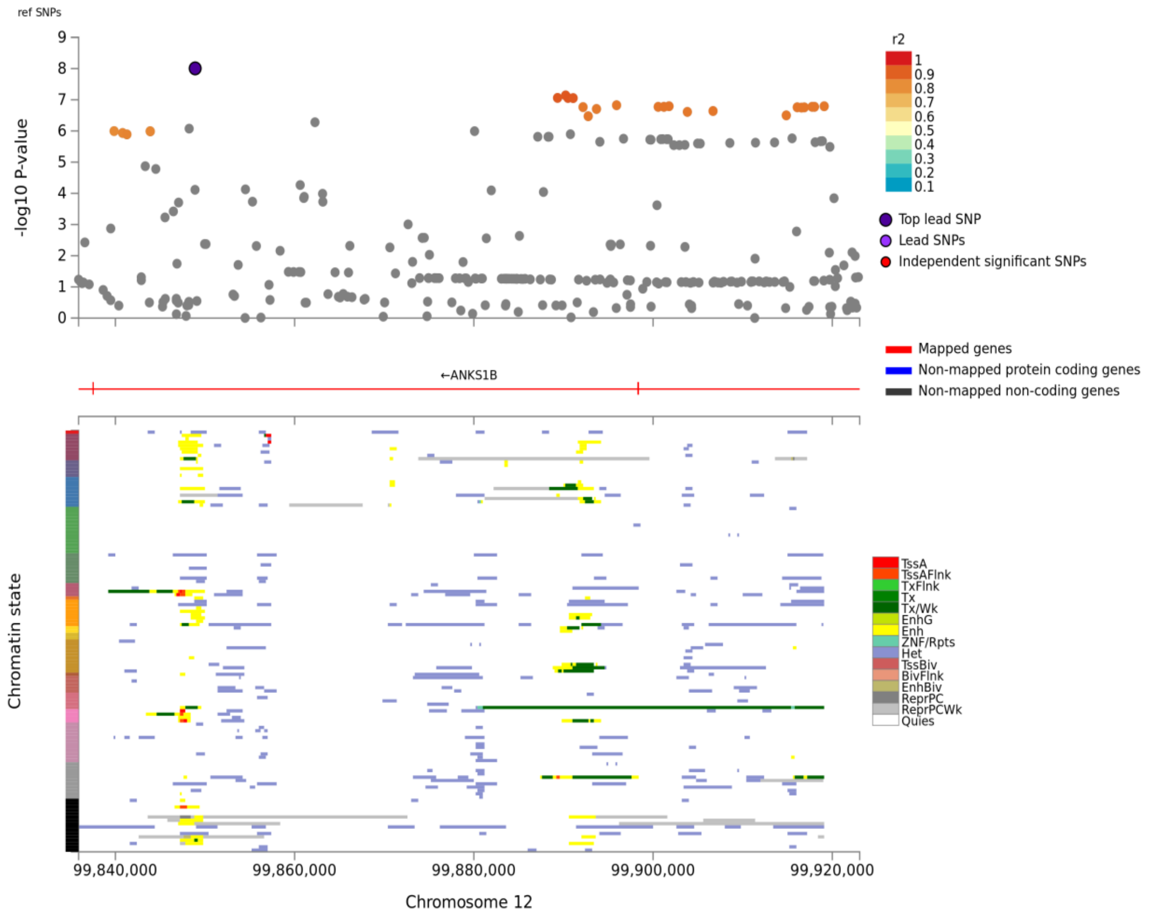


**Supplementary Figure 2** The 15-core chromatin state for the *ANKS1B* rs2133896 locus. The top lead SNP was in the enhancer region (yellow). TssA: Active TSS; TssAFlnk: Flanking Active TSS; TxFlnk: Transcript at gene 5’ and 3’; Tx: Strong transcription; TxWk: Weak transcription; EnhG: Genic enhancers; Enh: Enhancers; ZNF/Rpts: ZNF genes & repeats; Het: Heterochromatin; TssBiv: Bivalent/Poised TSS; BivFlnk: Flanking Bivalent TSS/Enh; EnhBiv: Bivalent Enhancer; ReprPC: Repressed PolyComb; ReprPCWk: Weak Repressed PolyComb; Quies: Quiescent/Low.

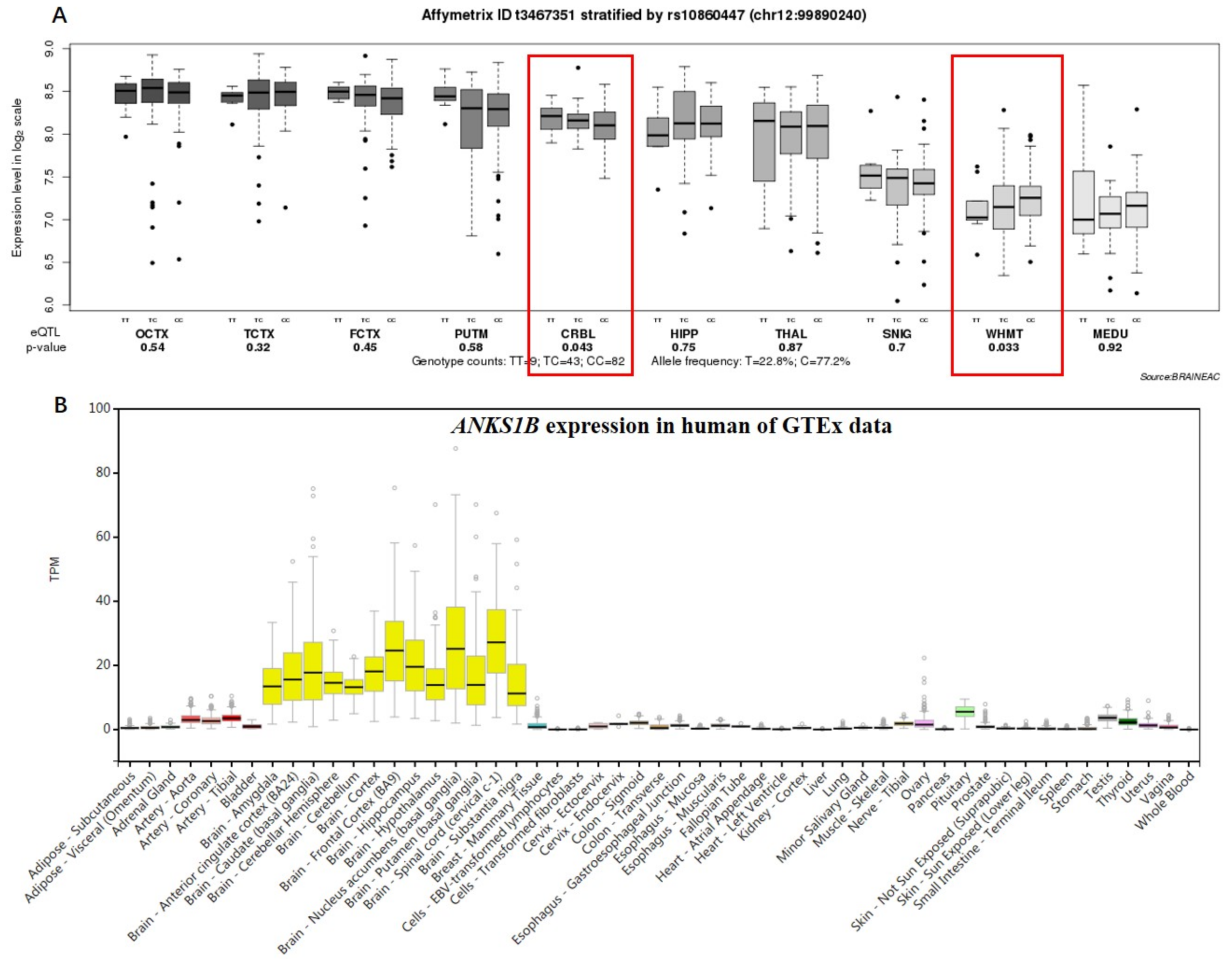


**Supplemental Figure 3. Stratified expression of *ANKS1B* by rs10860447 and *ANKS1B* gene expression profile. (A)** Effect of *ANKS1B* rs10860447 (LD-proxy with rs2133896 $r^2 = 0.902$) on gene expression in brain using BRAINEAC data. C allele (linked with rs2133896 T allele) had higher *ANKS1B* expression in white matter and lower expression in cerebellar cortex (P<0.05 but could not be corrected by multiple testing). CRBL, cerebellar cortex; FCTX, frontal cortex; HIPP, hippocampus; MEDU, medulla (specifically inferior olivary nucleus); OCTX, occipital cortex (specifically primary visual cortex); PUTM, putamen; SNIG, substantia nigra; TCTX, temporal cortex; THAL, thalamus; WHMT, intralobular white matter. (B) *ANKS1B* is specifically expressed in brain (GTEx data).

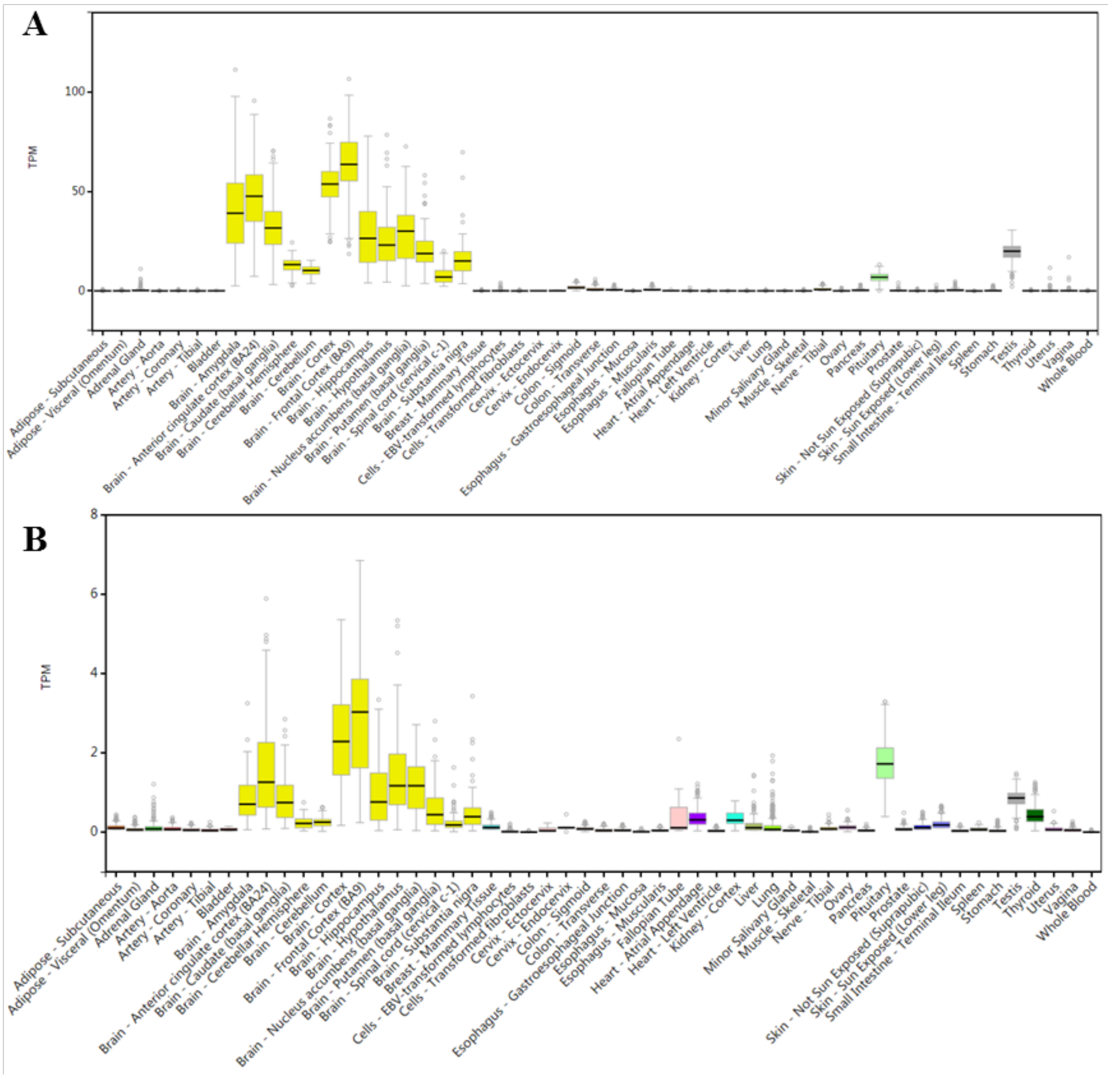


**Supplemental Figure 4.** GTEx gene expression profiles for *CTNNA2* (A) and *AGBL4* (B).

**Table** S1. Top 10 genes for gene-level analysis using MAGMA. Genes (in bold) passed multiple corrections. $p<0.05/18{,}225=2.74\times10^{-6}$.

| GENE | CHR | START | STOP | NSNPS | NPARAM | N | ZSTAT | P |
|---|---|---|---|---|---|---|---|---|
| **ADH1A** | **4** | **100192523** | **100217185** | **47** | **6** | **6155** | **5.5834** | **$1.18\times10^{-8}$** |
| **ADH1B** | **4** | **100222527** | **100247572** | **49** | **8** | **6155** | **5.5628** | **$1.33\times10^{-8}$** |
| **ADH6** | **4** | **100118795** | **100145403** | **34** | **6** | **6155** | **4.6742** | **$1.48\times10^{-6}$** |
| ANKS1B | 12 | 99123569 | 100383512 | 2103 | 94 | 6155 | 4.3931 | $5.59\times10^{-6}$ |
| MYO1H | 12 | 109821439 | 109891176 | 276 | 25 | 6155 | 4.3371 | $7.22\times10^{-6}$ |
| KCTD10 | 12 | 109881460 | 109920165 | 166 | 20 | 6155 | 4.0535 | $2.52\times10^{-5}$ |
| C22orf34 | 22 | 50008290 | 50056190 | 138 | 18 | 6155 | 3.9218 | $4.39\times10^{-5}$ |
| DYNLT1 | 6 | 159052506 | 159070804 | 30 | 5 | 6155 | 3.9202 | $4.42\times10^{-5}$ |
| CTNNA2 | 2 | 79735060 | 80880993 | 2495 | 167 | 6155 | 3.8815 | $5.19\times10^{-5}$ |
| ASPDH | 19 | 51009857 | 51023689 | 27 | 10 | 6155 | 3.8112 | $6.91\times10^{-5}$ |

**Table S2. Top 10 KEGG pathways for gene-set analysis using MAGMA. Pathways (in bold) passed multiple corrections. $p<0.05/221=2.26\times10^{-4}$.**

| Pathway Name | NGENES | BETA | BETA | SE | P |
|---|---|---|---|---|---|
| **Wnt signaling pathway** | **26** | **0.6377** | **0.02407** | **0.1625** | **$4.37\times10^{-5}$** |
| **Basal cell carcinoma** | **16** | **0.6871** | **0.0203** | **0.1913** | **$1.65\times10^{-4}$** |
| Bacterial invasion of epithelial cells | 70 | 0.3682 | 0.0228 | 0.1333 | $2.88\times10^{-3}$ |
| Ascorbate and aldarate metabolism | 70 | 0.3430 | 0.0212 | 0.1263 | $3.31\times10^{-3}$ |
| Hedgehog signaling pathway | 63 | 0.3518 | 0.0206 | 0.1336 | $4.23\times10^{-3}$ |
| Histidine metabolism | 47 | 0.3322 | 0.0168 | 0.1360 | $7.28\times10^{-3}$ |
| Folate biosynthesis | 26 | 0.5257 | 0.0198 | 0.2210 | $8.68\times10^{-3}$ |
| Drug metabolism – cytochrome P450 | 121 | 0.2180 | 0.0177 | 0.1000 | 0.0146 |
| T cell receptor signaling pathway | 41 | 0.3161 | 0.0150 | 0.1467 | 0.0156 |
| Selenoamino acid metabolism | 47 | 0.2752 | 0.0140 | 0.1342 | 0.0202 |

**Table S3. GRS association results for chr12 *ANKS1B* locus with addiction characteristics.**

| Addiction | Phenotype[a] | Estimate | Std.Error | t value | Pr(>\|t\|) |
|---|---|---|---|---|---|
| Alcohol | MAST | 2.726 | 10.812 | 0.252 | 0.801 |
| Alcohol | Age Onset | -19.916 | 18.414 | -1.082 | 0.281 |
| Alcohol | Dosage | -3.038 | 24.397 | -0.125 | 0.901 |
| MA | AgeOnset | 2.495 | 4.485 | 0.556 | 0.578 |
| MA | Frequency | 0.906 | 1.584 | 0.572 | 0.568 |
| MA | Dosage | -0.231 | 0.439 | -0.525 | 0.599 |
| Heroin | AgeOnset | 2.353 | 6.466 | 0.364 | 0.716 |
| **Heroin** | **Frequency** | **5.781** | **2.741** | **2.109** | **0.035** |
| Heroin | Dosage | 0.839 | 0.704 | 1.192 | 0.234 |

MA: methamphetamine; MAST: Michigan Alcoholism Screening Test.

[a] Dosage for heroin and MA was defined as g/per-time, dosage for alcohol was defined as 10 grams of alcohol (equivalent to 12.5 mL of pure alcohol).

**Table S4. Rs2133896 by group comparisons of mean FA for each of the 20 tracts.**

| White matter tract (20 tracts) | Heroin addicts (N=65) | | Healthy controls (N=69) | | F rs2133896×group (P) | F rs2133896 (P) | F group (P) |
|---|---|---|---|---|---|---|---|
| | G carriers (N=20) | TT genotype (N=45) | G carriers (N=36) | TT genotype (N=33) | | | |
| ATR.L | 0.41±0.02 | 0.41±0.02 | 0.41±0.02 | 0.41±0.02 | 0.065(0.799) | 0.094(0.760) | 1.127(0.290) |
| ATR.R | 0.41±0.01 | 0.41±0.02 | 0.40±0.02 | 0.41±0.01 | 0.069(0.793) | 2.370(0.126) | 4.921(0.028) |
| CST.L | 0.55±0.02 | 0.55±0.02 | 0.56±0.02 | 0.56±0.02 | 0.015(0.902) | 0.009(0.924) | 0.003(0.954) |
| CST.R | 0.55±0.02 | 0.56±0.02 | 0.55±0.02 | 0.55±0.02 | 0.214(0.644) | 0.032(0.858) | 0.237(0.627) |
| CCG.L | 0.52±0.03 | 0.52±0.04 | 0.50±0.03 | 0.51±0.04 | <0.001(0.988) | 0.004(0.952) | 10.145(0.002)* |
| CCG.R | 0.47±0.03 | 0.48±0.04 | 0.47±0.03 | 0.47±0.05 | 0.552(0.459) | 0.001(0.980) | 9.593(0.002)* |
| CHIP.L | 0.45±0.03 | 0.46±0.04 | 0.45±0.03 | 0.46±0.04 | 0.022(0.882) | 0.563(0.454) | 0.423(0.516) |
| CHIP.R | 0.43±0.03 | 0.43±0.04 | 0.43±0.04 | 0.42±0.04 | 0.107(0.745) | 0.585(0.446) | 1.289(0.258) |
| FMA | 0.65±0.02 | 0.65±0.02 | 0.65±0.02 | 0.65±0.02 | 1.335(0.250) | 0.116(0.734) | 0.543(0.462) |
| FMI | 0.51±0.01 | 0.51±0.02 | 0.50±0.02 | 0.51±0.02 | 0.010(0.920) | 1.586(0.210) | 3.005(0.085) |
| IFOF.L | 0.47±0.01 | 0.47±0.02 | 0.47±0.02 | 0.47±0.02 | 0.490(0.485) | 0.522(0.471) | 0.013(0.910) |
| IFOF.R | 0.46±0.02 | 0.46±0.02 | 0.46±0.02 | 0.46±0.02 | 0.797(0.374) | 0.174(0.677) | 0.434(0.511) |
| ILF.L | 0.43±0.01 | 0.44±0.02 | 0.44±0.02 | 0.44±0.02 | 2.630(0.107) | 2.730(0.101) | 0.435(0.511) |
| ILF.R | 0.43±0.02 | 0.44±0.02 | 0.44±0.02 | 0.43±0.03 | 3.349(0.069) | 1.547(0.216) | 0.485(0.487) |
| SLF.L | 0.43±0.01 | 0.44±0.02 | 0.44±0.02 | 0.44±0.02 | 4.735(0.031) | 2.779(0.098) | 0.810(0.370) |
| SLF.R | 0.43±0.02 | 0.44±0.02 | 0.44±0.02 | 0.44±0.02 | 10.433(0.002)* | 0.411(0.523) | 0.084(0.772) |
| UF.L | 0.45±0.03 | 0.44±0.03 | 0.45±0.03 | 0.44±0.02 | 0.159(0.691) | 0.070(0.791) | 0.251(0.617) |
| UF.R | 0.49±0.03 | 0.49±0.03 | 0.49±0.04 | 0.48±0.03 | 0.465(0.496) | 0.012(0.913) | 0.038(0.845) |
| SLFT.L | 0.47±0.04 | 0.46±0.03 | 0.46±0.03 | 0.46±0.03 | 0.506(0.478) | 0.704(0.403) | 1.507(0.222) |
| SLFT.R | 0.50±0.04 | 0.50±0.03 | 0.50±0.03 | 0.49±0.04 | 0.003(0.954) | 0.905(0.343) | 3.437(0.066) |

FA, fractional anisotropy; ATR, Anterior thalamic radiation; CST, Corticospinal tract; CCG, Cingulum (cingulated gyrus); CHIP, Cingulum (hippocampus); FMA, Forceps major; FMI, Forceps minor; IFOF, Inferior fronto-occipital fasciculus; ILF, Inferior longitudinal fasciculus; SLF, Superior longitudinal fasciculus; UF, Uncinate.fasciculus; SLFT, Superior longitudinal fasciculus (temporal part); L, left; R, Right. Data expressed as mean FA ± standard deviation. *P<0.05/20
